# Comprehensive 2D and 3D phenotypic characterization of human invasive lobular carcinoma cell lines

**DOI:** 10.1101/341313

**Authors:** Nilgun Tasdemir, Emily Bossart, Zheqi Li, Zhu Li, Kevin M. Levine, Britta M. Jacobsen, George C. Tseng, Nancy E. Davidson, Steffi Oesterreich

**Author notes:** Corresponding Author: Steffi Oesterreich, PhD, Women’s Cancer Research Center, UPMC Hillman Cancer Center, 204 Craft Avenue, Pittsburgh, PA 15213, USA. **Financial Support:** The work is in part funded by a Department of Defense Breakthrough Fellowship Award to NT (BC160764), Shear Family Foundation grant and Susan G. Komen Leadership grant to SO (SAC160073), Breast Cancer Research Foundation grants to NED and SO, and a grant (#4100070287) with Pennsylvania Department of Health. The Pennsylvania Department of Health specifically disclaims responsibility for any analyses, interpretations or conclusions. EB is supported by a National Institutes of Health (NIH) Ruth L. Kirschstein Award (1F31CA203055-01). ZL is supported by the University of Pittsburgh John S. Lazo Cancer Pharmacology Fellowship. KML is supported by an individual fellowship from the NIH/NCI (5F30CA203095) The project used the UPMC Hillman Cancer Center Biostatistics and Tissue and Research Pathology Services (TARPS) Cores, in part supported by P30CA047904.

## Abstract

Invasive lobular carcinoma (ILC) is the second most common subtype of breast cancer following invasive ductal carcinoma (IDC) and characterized by the loss of E-cadherin-mediated adherens junctions. Despite displaying unique histological and clinical features, ILC still remains a chronically understudied disease with limited knowledge on the available laboratory research models. To this end, herein we report a comprehensive 2D and 3D phenotypic characterization of four Estrogen Receptor-positive human ILC cell lines - MDA-MB-134, SUM44, MDA-MB-330 and BCK4. Compared to the IDC cell lines MCF7, T47D and MDA-MB-231, ultra-low attachment culture conditions revealed a remarkable anchorage-independence ability that was unique to the ILC cells, a feature not evident in soft agar gels. 3D Collagen I and Matrigel culture indicated a generally loose morphology for the ILC cell lines, which exhibited differing preferences for adhesion to ECM proteins in 2D. Furthermore, ILC cells had limited migration and invasion ability in wound-scratch and transwell assays with the exception of haptotaxis to Collagen I. Transcriptional comparison of the cell lines confirmed the decreased cell proliferation and E-cadherin-mediated intercellular junctions in ILC, while uncovering the induction of novel pathways related to cyclic nucleotide phosphodiesterase activity, ion channels, drug metabolism and alternative cell adhesion molecules such as N-cadherin, some of which were also differentially regulated in ILC versus IDC tumors. Altogether, these studies will serve as an invaluable resource for the breast cancer research community and facilitate further functional discoveries towards understanding ILC, identifying novel drug targets and ultimately improving the outcome of patients with ILC.

**Authors’ Contributions:** ***Conception and design:*** N. Tasdemir, NE. Davidson, S. Oesterreich

***Development of methodology:*** N. Tasdemir, L. Zhu, GC. Tseng, S. Oesterreich

***Acquisition of data (performed experiments, processed data, etc.):*** N. Tasdemir, E. Bossart, Z. Li, Z. Li

***Analysis and interpretation of data (e.g. biological interpretation, statistical analysis, computational analysis):*** N. Tasdemir, Z. Li, KM. Levine, NE. Davidson, S. Oesterreich

***Writing, review and/or revision of the manuscript:*** N. Tasdemir, Z. Li, KM. Levine, BM. Jacobson, GC. Tseng, NE. Davidson, S. Oesterreich

***Study supervision:*** NE. Davidson and S. Oesterreich

## Introduction

Invasive lobular carcinoma (ILC) is the second most common type of breast cancer following invasive ductal carcinoma (IDC), accounting for 10-15% of all cases (1). At an annual number of ~25-38,000, which is higher than ovarian cancer or melanoma, ILC is the 6^th^ most common cancer among women in US (2). Histologically IDC tumors form palpable masses or lumps, while ILCs grow as small, dyscohesive cells in a single-file pattern (1,3). This unique growth pattern makes mammographic detection and surgical removal of ILC difficult, complicating breast conservation (3). In addition, compared to IDCs, ILCs present more frequently as multi-centric and bilateral and with metastases to ovaries, peritoneum and gastrointestinal tract (1,4). Paradoxically, while patients with ILC display favorable prognostic and predictive factors (Estrogen Receptor [ER]-positive, Progesterone Receptor-[PR] positive, HER2-negative, low Ki67 index) and are mostly treated with endocrine therapy, they exhibit more long-term recurrences compared to patients with IDC, indicative of endocrine resistance (4,5).

Despite its distinctive histological and clinical features, ILC has remained a gravely understudied subtype of breast cancer. The most characteristic feature of ILC is the lack of E-cadherin-mediated adherens junctions, thought to be largely responsible for its single-file growth pattern (6). This hallmark E-cadherin loss, found in 95% of all ILC tumors versus in only 7% of IDCs, occurs through truncating mutations and loss-of-heterozygosity (6-8). Our knowledge of ILC as a unique subtype of breast cancer is only recently emerging with comprehensive reports from big consortia such as The Cancer Genome Atlas (TCGA) (7) and Rational Therapy for Breast Cancer (RATHER) (9). Multi-omics profiling of human tumors has begun to reveal candidate disease drivers such as HER2, HER3, FOXA1 and PIK3CA mutations, PTEN loss and ESR1 amplifications, events more frequently observed in ILC compared to IDC (7,9,10). However, the functional validation of these potential drivers is hindered by the availability of few ER-positive human ILC cell lines for use in the laboratory and limited knowledge on their biological phenotypes. Thus there is urgent need to develop additional cell line models, as well as thoroughly characterizing the cellular behaviors of the existing ones.

Our laboratory has recently reported the first profiling of ER function and endocrine response in ER-positive human ILC cell lines (11). Here we go one step beyond and characterize their growth and morphologies in 3D environments such as in ultra-low attachment (ULA) culture (12), soft agar (13), and within/on top of ECM proteins (14,15), as well as their adhesion properties in 2D (16). Using IDC cell lines for comparison, we probe their migration potential in response to both soluble attractants in chemotaxis assays (17) and to substrate bound ECM proteins in haptotaxis assays (18). In addition, we report on their abilities to invade Collagen I and Matrigel, as well as assessing their use of amoeboid invasion in non-cross-linked Collagen I gels (19,20). Comparison of transcriptional profiling data of ER-positive human ILC and IDC cell lines identified a number of clinically relevant genes and pathways that provide important insights into the sub-type specific gene expression programs likely responsible for their divergent biological phenotypes. Combined, our studies serve as invaluable resource for modeling ILC in the laboratory and pave the way for a promising direction of research for ILC biology towards new discoveries.

## Materials and Methods

### Cell culture

MDA-MB-134-VI (MDA-MB-134), MDA-MB-330, MCF-7, T47D and MDA-MB-231 were obtained from the American Type Culture Collection. SUM44PE (SUM44) was purchased from Asterand and BCK4 was kindly provided by Britta Jacobsen, University of Colorado Anschutz, CO. Cell lines were maintained in the following media (Life Technologies) with 10% FBS: MDA-MB-134 and MDA-MB-330 in 1:1 DMEM:L-15, MCF7 and MDA-MB-231 in DMEM, T47D in RPMI, BCK4 in MEM with non-essential aminoacids (Life Technologies) and insulin (Sigma-Aldrich). SUM44 was maintained as described (11) in DMEM-F12 with 2% charcoal stripped serum and supplements. Cell lines were routinely tested to be mycoplasma free, authenticated by the University of Arizona Genetics Core by Short Tandem Repeat DNA profiling and kept in continuous culture for <6 months. PIK3CA plasmids in pBABE (Addgene) and PTEN shRNAs in pMLPE (a kind gift from Scott W. Lowe, Memorial Sloan Kettering Cancer Center, NY) were packaged as previously described (21) and cells were selected with 1µg/ml puromycin (Life Technologies). shRNA sequences are provided in Supplementary Table 1.

### Immunoassays and trichrome staining

Western blots were performed as previously described (22) using 5% milk powder for blocking and developed using ECL (Sigma-Aldrich). For immunofluorescence, cells grown on coverslips were fixed with 4% paraformaldehyde, permeabilized and blocked with 5% Normal Goat Serum. Coverslips were incubated with antibodies, washed and mounted using DAPI- containing media (Thermo Fisher Scientific). Slides were imaged using a Nikon A1 advanced confocal system. Details of the antibodies used are included in Supplementary Table 1. Masson’s Trichrome (Sigma-Aldrich) staining was performed on a tissue microarray using manufacturer’s protocol without the acetic acid step after the aniline blue.

### Anchorage-independence and stemness assays

For 2D and ULA growth assays, ILC (15,000/96-well; 300,000/6-well) and IDC (5,000/96-well; 100,000/6-well) cells were seeded in regular (Thermo Fisher Scientific) or ULA (Corning Life Sciences) 96-well plates and assayed using CellTiter-Glo (Promega) on a Promega GloMax plate reader. Soft agar assays with ILC (50,000/plate) and IDC (10,000/plate) cells were performed in 35-mm plates (Thermo Fisher Scientific) as previously described (13,23). For mammosphere assays, ILC (60,000/well) and IDC (20,000/well) cells were seeded in 6-well ULA plates (Corning Life Sciences) as previously described (24) in 1:1 DMEM/Ham’s F-12 media with 20 ng/mL bFGF (BD Biosciences), 20 ng/mL EGF (BD Biosciences), B27 (Gibco), 2.5 mL Penicillin/Streptomycin, and 4 µg/mL Heparin (Sigma-Aldrich). All images were taken on an Olympus IX83 inverted microscope. For stem cell expression experiments, cells were stained with the indicated antibodies and analyzed on an LSRII flow cytometer (BD Biosciences). Gating was performed using the BD FACS Diva Software and isotype antibody stainings. Details of the antibodies used are included in Supplementary Table 1.

### 3D ECM assays

ILC (15,000/well) and IDC (5,000) cells were embedded in rat-tail Collagen I (Corning Life Sciences) at 4mg/ml in 24-well plates following manufacturer’s recommendations. For Matrigel (BD Biosciences) assays, ILC (5,000/well) and IDC (4,000/well) cells were seeded into single wells of 8-well LabTek Chamber Slides (Thermo Fisher Scientific) as previously described (14). Colonies were imaged on an Olympus IX83 inverted microscope.

### ECM adhesion assays

ILC (100,000-200,000/well) and IDC (50,000-10,000/well) cells were seeded into 96-well plates with the indicated coatings (Corning Life Sciences) following detachment with PBS containing 2mM EDTA. After incubation, plates were imaged on an Olympus IX83 inverted microscope, washed twice with PBS and quantified using FluoReporter dsDNA kit (Life Technologies) following manufacturer’s protocol.

### Migration and invasion assays

Wound-scratch assays were performed as previously described (25,26) using the IncuCyte Zoom Live Cell Imaging System (Essen Bioscience). PMA (Sigma-Aldrich) was used at 100 nM. For transwell experiments, cells were serum starved overnight, plated into 8 µm inserts (Thermo Fisher Scientific and Cell Biolabs) and quantified using Crystal violet following manufacturer’s protocol. Images of inserts were taken on an Olympus SZX16 dissecting microscope.

### Differential gene expression, pathway and survival analyses

RNA-Sequencing data of the cell lines was obtained from Marcotte et al. (27). The R package DESeq2 was used for differential expression analysis and pathway enrichment analysis was implemented using KEGG, BIOCARTA, REACTOME and KEGG databases as previously described (22). Survival analysis was performed using the METABRIC dataset (28) as previously described (22).

### Statistical analysis

Data analysis was performed using GraphPad Prism. Data is presented as mean +/- standard deviation. Statistical tests used for each figure are indicated in the respective figure legends.

### Supplementary methods

Detailed methods are described in Supplementary Text.

## Results

### Hormone receptor and adherens junction status

In this study, we focused on four ER-positive human ILC cell lines - MDA-MB-134, SUM44, MDA-MB-330 and BCK4 and utilized the IDC cell lines MCF7, T47D and MDA-MB-231 for comparative studies. As seen in **Fig. 1A**, Western blotting confirmed ER expression in all cell lines, except for the ER-negative MDA-MB-231 cells (29). Of note, although there are conflicting reports on the ER status of MDA-MB-330 cells (29,30), they displayed abundant ER levels in our hands. Only BCK4 and T47D had detectable PR expression at these exposure conditions. As expected, E-cadherin was absent from MDA-MB-134, SUM44, which harbor *CDH1* truncating mutations (30), from BCK4 cells, as well as from MDA-MB-231 cells, in which the *CDH1* promoter is hypermethylated (31).

**Figure 1.**
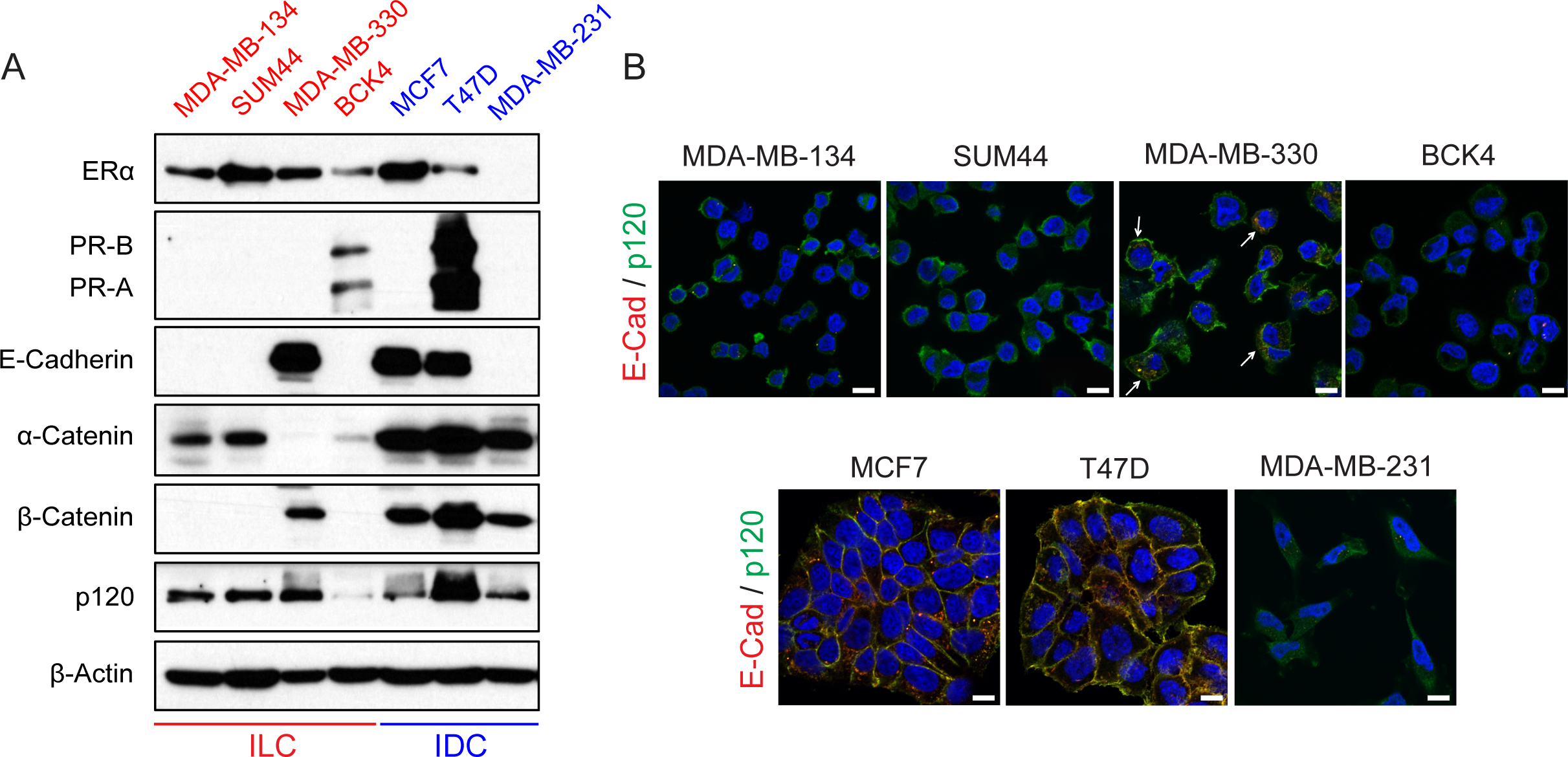
Characteristics of the human ILC and IDC cell lines used in the study. **A.** Western blotting using the indicated antibodies on whole cell lysates from ILC (left; red) and IDC (right; blue) cell lines. β-Actin was used as a loading control. **B.** Merged images of co-immunofluorescence staining for E-cadherin (red) and p120 (green) in ILC (top) and IDC (bottom) cell lines. DAPI (blue) was used for counterstaining to mark nuclei. Arrows indicate cytoplasmic co-localization of E-cadherin and p120 in MDA-MB-330 cells. Scale bar: 10 µm.

Consistent with the down-regulation and/or mislocalization of other junction components in the absence of E-cadherin (32,33), β-catenin expression was absent from MDA-MB-134, SUM44 and BCK4 cells – a result different from that in MDA-MB-231 cells, which retained β-catenin expression without E-cadherin. Interestingly, MDA-MB-330 cells, which harbor a bi-allelic, truncating *CTNNA1* mutation (30), still expressed E-cadherin and β-catenin in the absence of α-catenin. In contrast, p120-catenin (p120) was detected in all cell lines with the weakest expression in BCK4 cells, which also exhibited lower α-catenin levels. Co-immunofluorescence staining confirmed the absence of functional E-cadherin in ILC cell lines, which was mislocalized to the cytoplasm in MDA-MB-330 cells (**Fig. 1B**; top). Similarly, p120 was also largely cytoplasmic in ILC cell lines and in MDA-MB-231 cells, unlike its normal membranous co-localization with E-cadherin in IDC cells (**Fig. 1B**; bottom). Collectively, these data confirm the absence of functional adherens junctions in ILC cells.

### Anchorage-independence ability

Next we assessed the anchorage-independence ability of ILC cell lines by growing them in ULA conditions, which forces them into a suspension culture (12). ILC cell lines exhibited a dyscohesive, scattered morphology in 2D plates consistent with their lack of adherens junctions, while growing as large floating clusters in ULA plates (**Fig. 2A**; top). In contrast, MCF7 and T47D cells were more cohesive in 2D and formed tight spheres in ULA (**Fig. 2A**; bottom). Despite having overall slower proliferation rates compared to the IDC cells, the ILC cell lines had a remarkable ability to grow equally well in 2D and ULA plates, with BCK4 showing the least robust ULA phenotype (**Fig. 2B)**. Importantly, this anchorage-independence was unique to the ILC cells, as the IDC cell lines had much poorer growth in ULA versus 2D culture. Interestingly, while MDA-MB-231 cells with no adherens junctions displayed a loose ULA morphology more similar to ILC than IDC cells, they had the poorest ULA growth of all the IDC cell lines.

**Figure 2.**
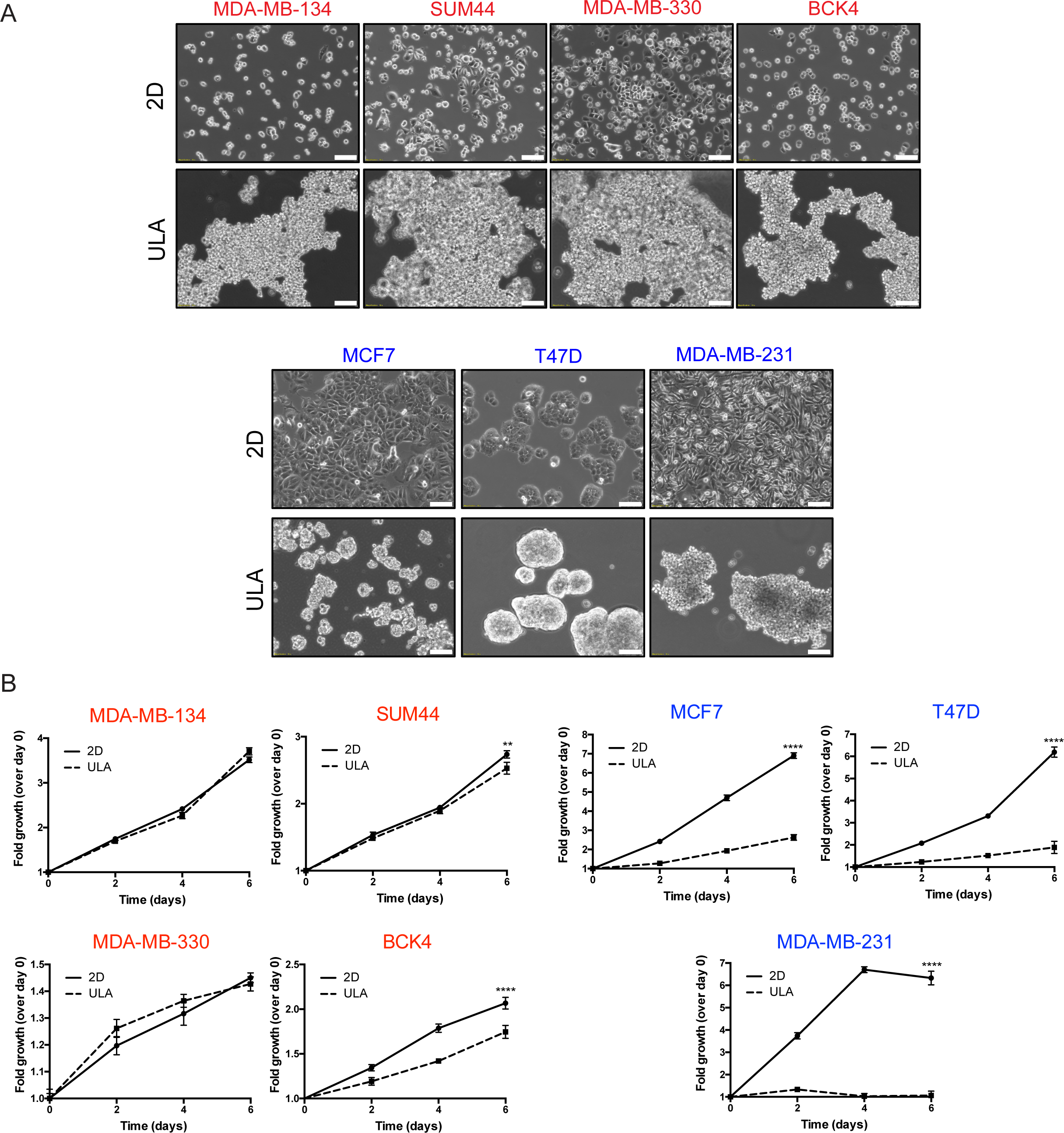
Human ILC cell lines exhibit superior growth in ULA culture than human IDC cell lines. **A.** Phase contrast light microscopy images of ILC (red; top) and IDC (blue; bottom) cell lines in 6-well 2D and ULA plates 4 days post plating. Scale bar: 100 µm. **B.** Relative growth curves showing fold growth normalized to day 0 at each time point over 6 days for ILC (red; left) and IDC (blue; right). Graphs show representative data from three experiments (n=6). p-values are two-way ANOVA comparison of 2D and ULA. * p ≤ 0.05; **** p ≤ 0.0001.

Given the superior anchorage-independent growth of ILC cell lines compared to IDC, we also assessed their ability to form mammospheres, which are similarly grown in ULA plates but with a selective media that enriches for stem-like cells (24). Assessment of stemness in the ILC cell lines was also of interest to us given the higher expression of stem cell markers such as ALDH1A1 in ILC versus IDC tumors (34,35). Despite their robust growth in ULA conditions, ILC cell lines formed poorly defined, loose mammospheres that were difficult to quantify, unlike the tighter MCF7 and T47D spheres (**Supplementary Fig. S1A**). Flow cytometric analysis of stem cell markers similarly did not identify a putative CD24^low^/CD44^high^ or CD49f^high^/EPCAM^low^ stem cell population in the ILC cell lines (**Supplementary Fig. S1B-C)**. Although such a population was present in MDA-MB-231 cells, this cell line did not form mammospheres as robustly as MCF7 and T47D cells. Consistent with previous literature, these results indicate poor mammosphere formation in cells with disrupted adherens junctions and a discordance between stem cell expression and mammosphere formation ability (36).

Another form of anchorage-independence is the ability to grow in suspension in soft agar gels (13). In general, ILC cell lines exhibited limited, dyscohesive growth in this semi-solid medium, with BCK4 cells forming the smallest colonies (**Fig. 3A;** top). The ILC growth was similar to the growth of MCF7 and T47D cells, the latter displaying a tighter morphology. As expected, MDA-MB-231 cells formed the most robust soft agar colonies, serving as a positive control. Altogether, these assays indicate that ILC cells exhibit a unique anchorage-independence ability in ULA conditions, a phenotype not replicated in soft agar or mammosphere culture.

**Figure 3.**
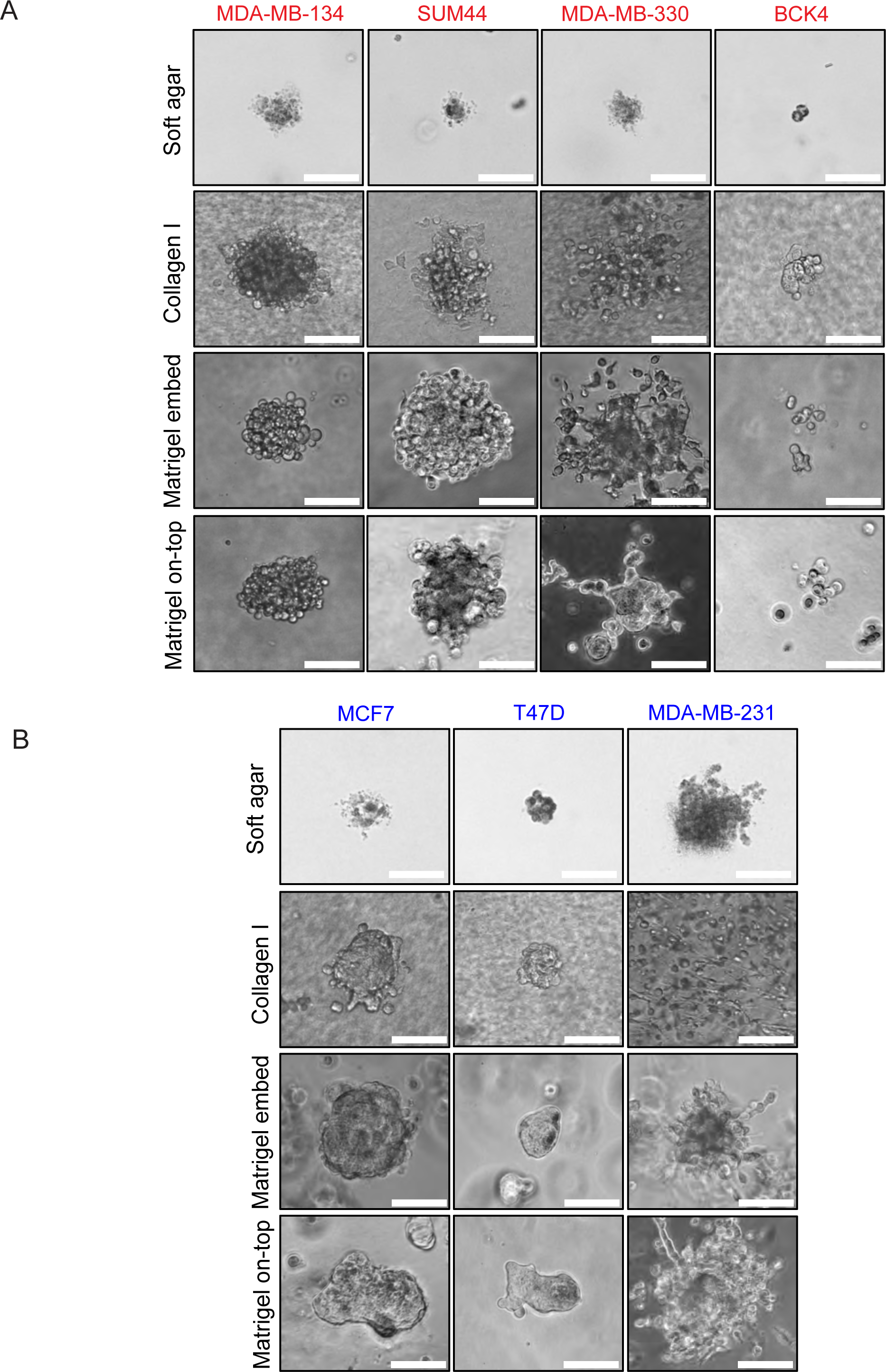
ILC and IDC cell lines exhibit varying morphologies in 3D culture. Phase contrast light microscopy images of (**A**) ILC (red) and (**B**) IDC (blue) cell lines in soft agar (4 weeks post plating), Collagen I, Matrigel embedded and Matrigel on-top culture (ILC: 3 weeks post plating; IDC: 1 week post plating). Scale bar: 100 µm.

### 3D ECM growth and cell-matrix interactions

In tissues, ILC tumors grow as a single-file of cells within a dense layer of stroma rich in ECM (1,6). This phenomenon can be visualized by staining human tumors with Masson’s Trichrome, which clearly demonstrates higher levels of collagen fibers in ILC compared to IDC (**Supplementary Fig. S2**). However, the 3D ECM growth of human ILC cell lines has not previously been systematically analyzed. Therefore, we first embedded ILC cell lines in thick Collagen I gels, where MDA-MB-134 and SUM44 cells exhibited the most robust growth, while MDA-MB-330 cells displayed a looser morphology and BCK4 formed the smallest colonies (**Fig. 3A**). Similar results were obtained when the cells were either embedded within or cultured on top of Matrigel, displaying a “grape-like” morphology previously described for cells with poor cell-cell adhesion (37). In contrast, MCF7 and T47D cells formed very tight colonies in all ECM environments (**Fig. 3B**). Interestingly, MDA-MB-330 cells exhibited protrusive structures in Matrigel culture (**Fig. 3A**), which are more characteristic of MDA-MB-231 cells with a “stellate” morphology and known invasive potential (**Fig. 3B**) (37).

In addition to growing cells within 3D gels, we also assayed the adhesion of cells to ECM proteins in 2D to gain a deeper understanding of their cell-matrix interactions (16). To this end, we seeded ILC and IDC cell lines onto plates coated with Collagen I, Collagen IV, Fibronectin, Laminin or Matrigel in serum free media. We also utilized uncoated plates for comparison and bovine serum albumin (BSA) coated plates as negative control for background adhesion levels. While a 2-hour incubation indicated a low level of overall binding in ILC cell lines, especially in MDA-MB-134 and SUM44 cells (**Supplementary Fig S3A)**, a 16 hour-incubation resulted in more efficient binding and varying cell morphologies on different matrices (**Fig. 4A** and **Supplementary Fig. S3B**). The ECM protein most preferred for binding in general was Collagen I (**Fig. 4B**), on which most cells displayed prominent adhesive protrusions (**Fig. 4A**), followed by Collagen IV and Matrigel.

**Figure 4.**
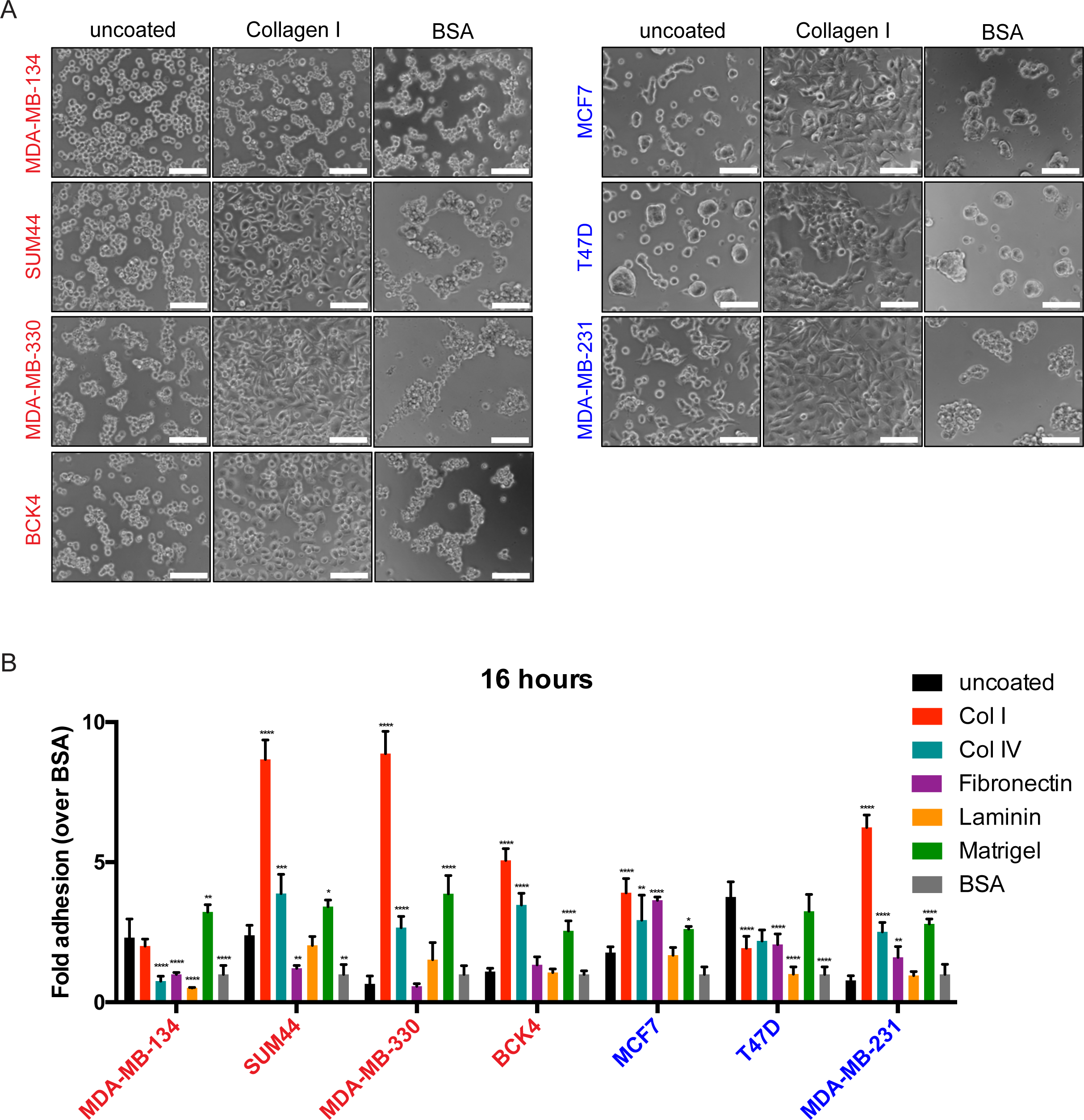
Human ILC cell lines have differing preferences for adhesion to ECM proteins. **A.** Phase contrast light microscopy images of ILC (red; left) and IDC (blue; right) cell lines in uncoated, Collagen I or BSA-coated plates 16 hours post-plating. Scale bar: 100 µm. **B.** Fold adhesion (normalized to BSA) of ILC (red) and IDC (blue) cell lines 16 hours post-plating. Graphs show representative data from two experiments (n=4). p-values are from ordinary one-way ANOVA with Dunnett’s multiple comparison test to uncoated conditions for each cell line. *p ≤0.05; **p ≤0.01; ****p ≤0.0001.

Of the ILC cell lines analyzed, MDA-MB-134 cells displayed a unique matrix interaction profile, with a less overall adhesion to ECM proteins than to uncoated plates and no visible adhesive protrusions on any matrix. Interestingly, unsupervised clustering of the ILC and IDC cell lines using publicly available transcript profiling data (27) showed that MDA-MB-134 cells clustered separately from the other ILC cell lines, displaying a unique expression pattern of genes encoding both integrins (**Supplementary Fig. S3C)** and matrix metalloproteinases (MMPs) (**Supplementary Fig. S3D)**, which are well known mediators of cell-matrix adhesion (16). Combined, these data indicate that ILC cell lines have differing morphologies in 3D ECM gels and divergent adhesive properties on matrix proteins.

### Migration and invasion potential

Next we assessed cell migration employing the commonly used wound-scratch assay, in which a gap (“a wound”) is introduced into the middle of a monolayer to induce directional movement of cells from the wound edges (17,25). Using the IncuCyte live-cell imaging system and capturing images of cells every 4 hours, we observed very limited basal migration in the ILC cell lines (**Fig. 5A**; left). This was in stark contrast to the IDC cells (**Fig. 5A**; right), which completely closed the wound in as early as 24 hours (MDA-MB-231). To exogenously induce cell migration, we treated the cells with phorbol myristate acetate (PMA), which activates the PKC pathway and downstream actin cytoskeleton reorganization (38). PMA treatment clearly triggered migratory protrusions at the edges of the wound in MCF7 cells (**Fig. 5B;** right) and substantially increased their migration rate (**Fig. 5C;** right). In contrast, however, despite inducing protrusions in the otherwise-round ILC cell lines (**Fig. 5B;** left), PMA had limited effect on their movement (**Fig. 5C**; left). While the strongest PMA effect was in BCK4, this cell line still failed to close the wound after 72 hours.

**Figure 5.**
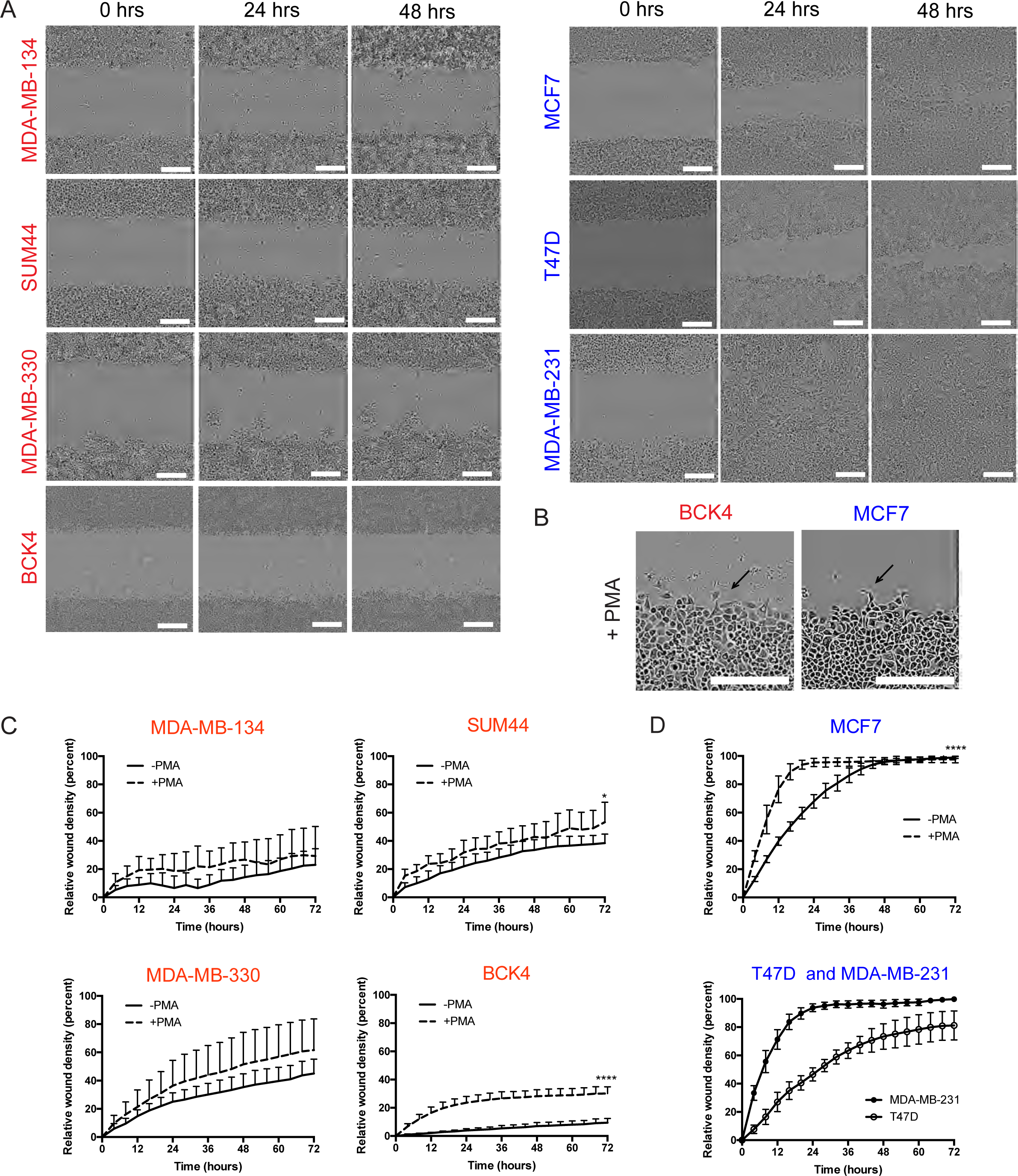
Human ILC cell exhibit limited migration ability in wound-scratch assays. **A.** Snapshots of the scratch wounds in ILC (red; left) and IDC (blue; right) cell lines at the indicated time points. Scale bar: 100 µm. **B.** Snapshots of the scratch wounds in BCK4 (red) and MCF7 (blue) cell lines treated with 100 nM PMA. Arrows indicate migratory protrusions at the wound edge. Scale bar: 300 µm. **C**. Relative wound densities over hour 0 in ILC (red; left) and IDC (blue; right) cell lines over time with and without PMA treatment. Graphs shows representative data from two-three experiments (n=6-8). p-values are from two-way ANOVA comparison of-PMA vs +PMA. * p ≤ 0.05; **** p ≤ 0.0001.

As an alternative to the wound-scratch assay, we next utilized transwell Boyden chambers to assess migration and invasion (17). As expected, the highly migratory MDA-MB-231 cells exhibited substantial chemotaxis to FBS, while MCF7 and T47D cell were weakly migratory (**Fig. 6A**). However, the ILC cell lines exhibited very limited migration to FBS in this assay. Given the ECM-rich stroma of ILC tumors (1,6) (see **Supplementary Fig. S2**), we also assayed migration to substrate bound ECM in haptotaxis experiments (18), in which the undersides of the inserts were coated with a thin layer of Collagen I. Interestingly, SUM44 and MDA-MB-330 cells displayed abundant haptotaxis to Collagen I over BSA in this assay (**Fig. 6B**), despite no chemotaxis to FBS (see **Fig. 6A)**. This result was different from MCF7 and MDA-MB-231 cells, which exhibited Collagen I haptotaxis (**Fig. 6B**) but also chemotaxis to FBS (see **Fig. 6A**), highlighting the unique requirement of matrix only by ILC cells for migration. However, this finding did not extend to BCK4 or MDA-MB-134 cells, which did not migrate substantially in either assay (**Fig. 6A-B**), with the phenotype of the latter being consistent with its weak ECM adhesion (see **Fig. 4**).

**Figure 6.**
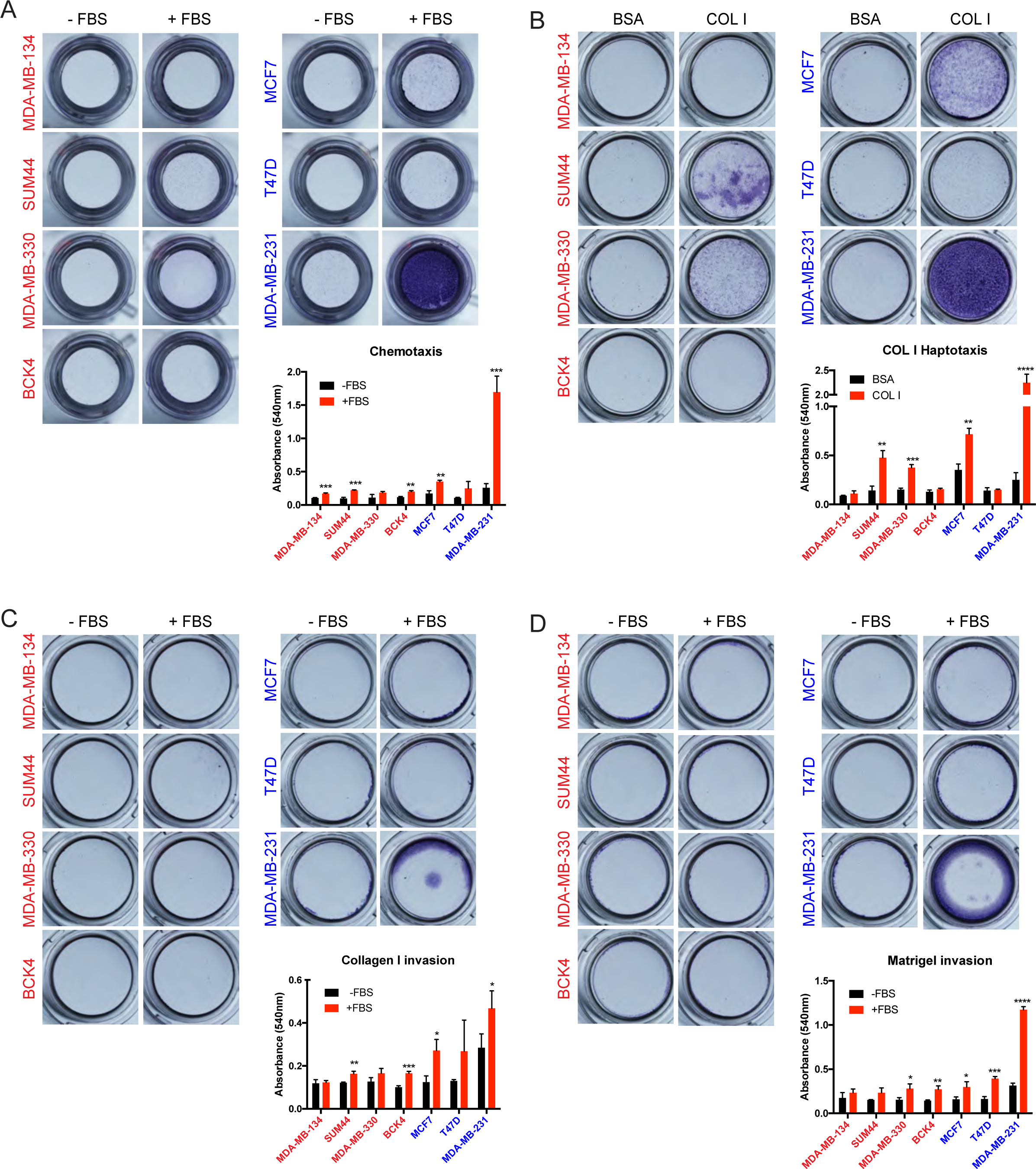
Human ILC cell lines exhibit limited migration and invasion towards FBS but SUM44 and MDA-MB-330 exhibit haptotaxis to Collagen I in transwell Boyden chamber assays. Images (top) and quantification (bottom) of crystal violet-stained inserts with ILC (red; left) and IDC (blue; right) cell lines from (**A**) Chemotaxis (**B**) Haptotaxis (**C**) Collagen I invasion and (**D**) Matrigel invasion assays towards the indicated attractants after 72 hours. Graphs show representative data from two independent experiments (n=3 biological replicates). p-values are from unpaired t-test. * p ≤ 0.05; ** p ≤ 0.01; *** p ≤ 0.001; **** p ≤ 0.0001.

Almost half of all human ILC tumors harbor activating, hotspot mutations in PIK3CA and 13% have PTEN loss due to inactivating mutations or deletions (7). These alterations are known to activate downstream Akt signaling, which can induce cell migration and invasion (39). While MCF7 and T47D both harbor PIK3CA mutations, human ILC cell lines are wild type for PIK3CA and PTEN (27). We therefore overexpressed PIK3CA mutants and knocked down PTEN in MDA-MB-134 cells to potentially augment their migration ability. Interestingly, compared to the respective controls, only the H1047R but not the E545K mutation activated downstream Akt signaling, along with all four PTEN shRNAs in MDA-MB-134 cells, which was different from the highly active MCF7 cells that harbor the endogenous E545K mutation (**Supplementary Fig. 4A**). However, when assayed in either transwell Boyden chambers (**Supplementary Fig. 4B-C**) or wound-scratch assays (**Supplementary Fig. 4D**), none of the tested alterations alone were sufficient to induce cell migration in MDA-MB-134 cells.

Using the transwell Boyden chambers, we also assayed the invasion capacity of human ILC cell lines. When plated in top chambers coated on the inside with either acid-extracted cross-linked Collagen I (**Fig. 6C**) or Matrigel (**Fig. 6D**), the highly invasive MDA-MB-231 cells were the only cell line that exhibited robust invasion; while MCF7 and T47D were weakly invasive. The ILC cell lines, however, had limited invasion of either Collagen I or Matrigel in response to FBS in bottom chambers. In addition to mesencyhmal invasion of cross-linked ECM proteins, cells can also exhibit amoeboid invasion by squeezing through the pores in non-cross-linked ECM (19,20). Given their morphological similarity to the small, round cells of melanoma and non-small cell lung cancer that utilize amoeboid invasion, we also assayed this type of invasion in ILC cell lines using transwell chambers coated with pepsin-extracted Collagen I. In contrast to the invasive MDA-MB-231 cells, however, ILC and IDC cell lines exhibited limited ameoboid invasion in this assay (**Supplementary Figure 5**). Altogether, these results suggest that ILC cell lines exhibit limited migration and invasion in traditional laboratory assays with the exception of haptotaxis to Collagen I.

### Transcriptional comparison of ILC and IDC cell lines and tumors

Our comprehensive analysis of the 2D and 3D phenotypes of human ILC cell lines clearly demonstrated unique biological properties. In order to delineate the gene expression programs that may underlie the divergent cellular phenotypes of the ILC and IDC cell lines, we performed transcriptional comparison analyses using publicly available data sets (27), which covered all of the cell lines used in this study except for BCK4. Importantly, of the ILC cell lines with available data, we only focused on MDA-MB-134 and SUM44 cells in order to capture the differential expression of E-cadherin in ILC versus IDC and therefore excluded MDA-MB-330 cells that do not harbor the hallmark *CDH1* mutation. In addition, ER-negative MDA-MB-231 cells were also excluded from the analyses to ensure comparison between cell lines belonging to the same molecular subtype (i.e. luminal), leaving MCF7 and T47D.

Despite the small number of cell lines analyzed, unsupervised hierarchical clustering of the ILC and IDC cells clustered MDA-MB-134 and SUM44 closer to each other and away from MCF7 and T47D (**Supplementary Fig. S6A**). Differential expression analysis, based on a fold-change cut-off value of 1.5 and false discovery rate (FDR) value of 0.05, identified 320 genes that were expressed higher in ILC versus IDC cell lines and 387 that were expressed lower (**Fig. 7A** and **Supplementary Table 2**). Pathway enrichment analysis on the differentially expressed genes (**Fig. 7B** and **Supplementary Table 3)** indicated upregulated transmembrane protein tyrosine kinase pathway in ILC cell lines, consisting of genes such as *FGFR1*, a known amplified oncogene in ILC (40). Additional pathways revealed from this analysis included ion channel activity, tyrosine metabolism, biological oxidation, cyclic nucleotide phosphodiesterase activity, drug metabolism cytochrome P450 and alternative cell adhesion, with genes such as *CDH2*. Conversely, the downregulated pathways confirmed the decreased intercellular junctions and proliferation in ILC versus IDC cell lines (6,30), as well as extending to further categories such as interferon signaling, amyloids, RNA Pol I transcription, extracellular structure and organization, and focal adhesions, with the last category mediating cell-matrix interactions (**Fig. 7C** and **Supplementary Table 3)**.

**Figure 7.**
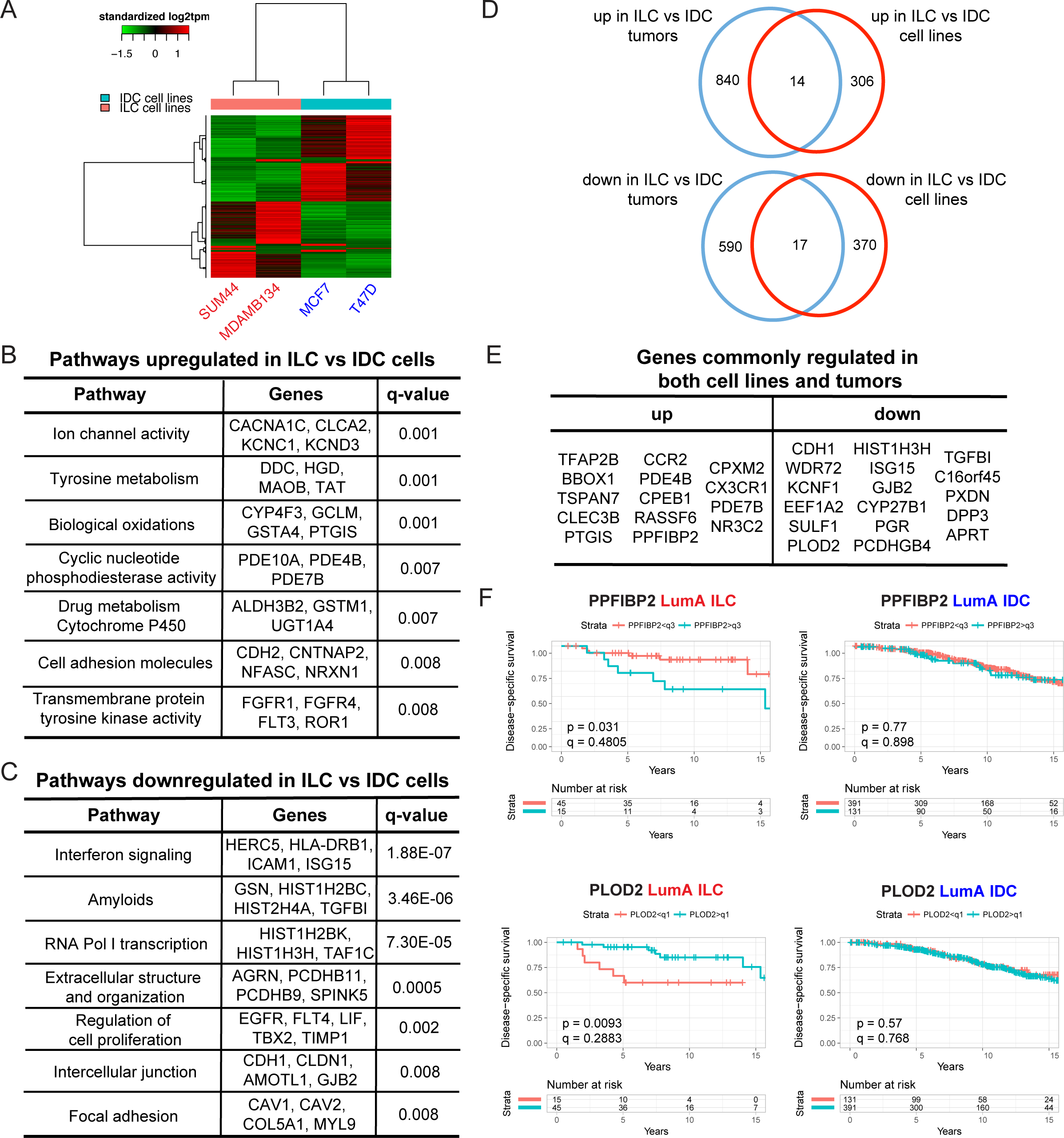
Transcriptional differences between human ILC and IDC cell lines and tumors. **A.** Supervised clustering heat map of ILC (red) and IDC (blue) cell lines using differentially expressed genes. **B-C.** Lists of representative (**B**) upregulated and (**C)** downregulated pathways and genes in ILC versus IDC cell lines. q-values are from Fisher’s exact test with FDR<0.05. **D**-**E**. Venn diagrams (**D**) and lists (**E**) of commonly upregulated and downregulated genes between ILC and IDC cell lines and tumors. **F.** Disease-specific survival curves for PPFIBP2 (top) and PLOD2 (bottom) in ER+, Luminal A ILC (red; left) and IDC (blue; right) patients in the METABRIC dataset. Patients were divided into two groups by PPFIBP2 (q3: third quadrant) and PLOD2 (q1: first quadrant) expression. p-values are from log-rank test. q-values were calculated using the Benjamini-Hochberg method to correct p-values for multiple comparisons testing within histology. The small ILC sample size (n=60) allows for limited statistical power in detecting survival differences after multiple testing correction.

We recently reported a transcriptomic comparison of ILC and IDC tumors from the TCGA (7) and METABRIC (28) cohorts (22). We therefore wished to determine to what extent the *in vitro* transcriptional differences from the cell lines correlated with their *in vivo* counterpart from the tumors. To this end, we initially performed a principal component (PCA) analysis of all the TCGA ER-positive, luminal A ILC (n=174) and IDC (n=774) tumors used in our original study (22) and the four cell lines used herein (MDA-MB-134, SUM44, MCF7, T47D), which somewhat separated the ILC and IDC tumors but clustered all the cell lines separately and away from the tumors (**Supplementary Fig. 6B**). Nevertheless, overlap of the differentially expressed genes between ILC and IDC tumors (22) and those between ILC and IDC cell lines (**Supplementary Table 2)** identified 14 upregulated and 17 downregulated genes, the latter including *CDH1* (**Fig. 7Dx-E**). The query of these genes in the METABRIC dataset (28) revealed that higher expression of *PPFIBP2*, as seen in cell lines and tumors from ILC versus IDC, was significantly associated with worse disease-specific survival in ILC but not in IDC (**Fig. 7F;** top). Similarly, lower expression of *PLOD2*, as seen in ILC versus IDC tumors and cell lines, exhibited a significant association with worse disease-specific survival in ILC but not in IDC (**Fig. 7F;** bottom). Collectively, these data highlight the sub-type specific, clinically relevant gene expression programs that may account for the divergent biological phenotypes between ILC and IDC cell lines and should be more deeply explored.

## Discussion

ILC is a special subtype of breast cancer with distinct histological and clinical features from the more common subtype IDC (1). Despite the clear need to expand our understanding of the unique biology of ILC, there are currently few laboratory models available for research and a huge gap in our knowledge on their biological properties beyond endocrine response (11). Our study is the first comprehensive report on the 2D and 3D phenotypic characterization of four ER-positive human ILC cell lines. Using a number of IDC cell lines for comparison, herein we profiled their 2D and 3D growth, matrix interactions, migratory and invasive properties and sub-type specific gene expression programs. Although a number of ER-negative ILC cell lines are available (30), we limited our focus to the ER-positive models given that approximately 90% of ILC tumors are of this molecular subtype (1,6). Interestingly, despite the controversial ER status of MDA-MB-330 cells (29,30), we included them in our studies since in our hands they exhibited abundant expression of ER, the functionality of which needs further investigation.

The unique anchorage-independence ability we observed in the human ILC cell lines suggests resistance to anoikis - detachment induced apoptosis -, and is in agreement with findings in mouse cell lines from a transgenic model of ILC (41). Given the well-accepted role of anchorage-independence in metastasis (42), our interesting result may have important clinical implications. While ILC and IDC tumors both progress to the stage of disseminated disease, patients with ILC present more often with long-term endocrine-resistant recurrences (1,4,5). The ability to survive in the absence of attachment to matrix may allow ILC cells to stay dormant in foreign ECM environments for extended periods of time prior to re-growth and colonization. Interestingly, our results indicated that this anchorage-independence ability was unique to the ILC cells in the ULA settings and not evident in soft agar or mammosphere culture, suggesting a context-dependent phenotype. Importantly, based on our data, testing of therapeutic agents for ILC in future studies in both 2D and ULA conditions might allow uncoupling of potential effects on cell proliferation versus metastasis.

Our 3D ECM experiments revealed generally loose, poorly defined colonies for the ILC cell lines, which is not surprising given their defect in adherens junctions (6). Nevertheless, different cell lines still displayed varying morphologies and abilities to grow in these settings. Such a divergent pattern was especially evident in the ECM adhesion experiments, where MDA-MB-134 generally exhibited a less preference for interacting with matrix proteins. Our *in silico* analysis revealed a putative list of integrins and MMP proteins that may account for this phenotype. Interestingly, none of the ILC cell lines analyzed assumed their native, *in vivo* single-file morphology within or on top of 3D ECM gels on 2D matrix coatings. Since the single-file pattern of the cancer cells in ILC tumors may provide important spatial and polarity cues, forcing ILC cell lines to grow in linear patterns using platforms such as micro-patterned ECM surfaces (43) may allow better modeling of ILC in the laboratory. Based on our data, the poor ECM adhesion of MDA-MB-134 cells makes them a less suitable choice for such future studies compared to the remaining ILC cell lines.

Given their dyscohesive morphology, we expected that ILC cell lines may exhibit single cell migration as opposed to the collective migration of cell lines such as MCF7. To our surprise, however, in traditional wound-scratch assays, ILC cells exhibited a remarkable inability to robustly migrate even with PKC or Akt activation. This result may be due to their lack of adherens junctions, which makes it difficult to grow them into a complete monolayer and likely prevents them from experiencing the same loss of cell polarity at the wound edge as IDC cells. In transwell Boyden chambers, ILC cell lines did not exhibit substantial migration or invasion except for haptotaxis of SUM44 and MDA-MB-330 cells to Collagen I, which was consistent with their matrix adhesion properties and highlights the need for incorporating ECM proteins into such assays. While our attempts at studying amoeboid invasion of ILC cells in non-crosslinked Collagen I did not reveal much movement towards FBS, there is a clear need for more sophisticated, alternative assays using micro-patterned surfaces and stromal cell types such as fibroblasts to generate physiologically-relevant confined spaces and ECM tracks (43,44).

Our transcriptional comparison of ILC and IDC cell lines confirmed previously known differences in E-cadherin-mediated cell-cell junctions, cell proliferation and expression of transmembrane receptor tyrosine kinases such as *FGFR1* (6,30,40). In addition, this analysis revealed a number of novel differences in pathways such as ion channel activity, drug metabolism cytochrome P450 and extracellular structure and organization. Interestingly, one pathway upregulated in ILC versus IDC cell lines was related to cell-cell adhesion and consisted of genes such as *CDH2*, which encodes N-cadherin. This pathway may provide alternative cell-cell communication in the absence of E-cadherin and may be indicative of a partial epithelial-to-mesenchymal transition in ILC, a result consistent with the recent RATHER report on ILC tumors (9). Collectively, the genes and pathways we identified may account for the divergent biological phenotypes of the ILC and IDC cell lines we observed throughout our studies.

Overlaying the gene expression data from ILC and IDC cell lines with that from tumors, we observed a completely separate clustering and very little overlap. This phenomenon has previously been reported for ovarian cancer (45) and is not surprising given the much higher complexity of *in vivo* settings compared to *in vitro* (15,46). Future gene expression profiling studies of the cell lines cultured on ECM proteins analyzed in our study and/or in the presence of stromal cell types such as fibroblasts should yield a better recapitulation of tumor transcriptional programs. Nevertheless, our analysis identified a number of differentially regulated genes that were common between the cell lines and tumors, some of which exhibited a significant correlation with disease-free survival of ILC but not IDC tumors. This approach helped generate a short list of clinically relevant genes that may be pertinent to the unique ILC biology.

*PPFIBP2*, also known as Liprin Beta 2, encodes a protein involved in the plasma membrane recruitment of leukocyte common antigen-related receptor (LAR) protein-tyrosine phosphatases, which regulate focal adhesions and mammary gland development (47). It is commonly fused to oncogenes such as RET in thyroid cancer and the hypo-methylation of its enhancer is associated with increased breast cancer risk (48). Since our data links *PPFIB2* to poor survival specifically in the ILC cohort, further functional interrogation of this understudied, candidate oncogenic driver may implicate it as a novel therapeutic target in ILC. PLOD2 encodes an enzyme involved in the hydroxylation of lysyl residues in collagen-like peptides and ECM remodeling (49). In contrast to its promotion of metastasis in lung adenocarcinoma (50), the association of *PLOD2* with better survival specifically in the ILC cohort suggests that decreased collagen crosslinking may create a microenvironment more permissive to growth and dissemination in ILC, underlining the importance of studying amoeboid migration and invasion.

In conclusion, our comprehensive characterization of the 2D and 3D phenotypes of ER-positive human ILC lines revealed important insights into the unique biology of ILC. With increasing interest in ILC in the laboratory and a growing list of candidate disease drivers from next-generation sequencing efforts, our study will serve as an invaluable resource for the breast cancer research community and as a platform to facilitate functional validation of potential therapeutic targets towards improving the clinical outcome of patients with ILC.

## Acknowledgments

The authors thank Dr. Jennifer Xavier for critical reading of the manuscript.

